# The intrinsically disordered protein glue of myelin: Linking AlphaFold2 predictions to experimental data

**DOI:** 10.1101/2022.09.13.507838

**Authors:** Oda C. Krokengen, Arne Raasakka, Petri Kursula

## Abstract

Numerous human proteins are either partially or fully classified as intrinsically disordered proteins (IDPs). Due to their properties, high-resolution structural information about IDPs is generally lacking. On the other hand, IDPs are known to adopt local ordered structures upon interactions with ligands, which could be *e.g*. other proteins or lipid membrane surfaces. While recent developments in protein structure prediction have been revolutionary, their impact on IDP research at high resolution remains limited. We took a specific example of two myelin-specific IDPs, the myelin basic protein (MBP) and the cytoplasmic domain of myelin protein zero (P0ct). Both of these IDPs are known to be crucial for normal nervous system development and function, and while they are disordered in solution, upon membrane binding, they partially fold into helices, being embedded into the lipid membrane. We carried out AlphaFold2 predictions of both proteins and analysed the models in light of previously published data related to solution structure and molecular interactions. We observe that the predicted models have helical segments that closely correspond to the characterised membrane-binding sites on both proteins. We furthermore analyse the fits of the models to SAXS data from the same IDPs. Artificial intelligence-based models of IDPs appear to be able to provide detailed information on the ligand-bound state of these proteins, instead of the form dominating free in solution. We further discuss the implications of the predictions for normal mammalian nervous system myelination and their relevance to understanding disease aspects of these IDPs.

## Introduction

The artificial intelligence/machine learning-based algorithms of protein structure prediction, most notably AlphaFold2 [1] and RoseTTAFold [2], have recently revolutionised structural biology. AlphaFold2 is trained on crystal structures from the Protein Data Bank, which suggests it will predict conformations that one might find in a protein crystal, and for many folded proteins, the predictions are essentially identical to the crystal structure – sometimes even allowing error detection in the experimental structure [3]. With the development of AlphaFold2, the structural coverage of all human protein residues has gone from 48% to 76%. Thus, the number of proteins with no structural coverage is now down to 29% [4]. It is, however, obvious that for intrinsically disordered proteins (IDPs) and flexible multidomain proteins with intrinsically disordered regions (IDRs), AlphaFold2 cannot predict a single accurate 3D structure – which in such cases does not even exist.

Considering the above, the fold predictions of AlphaFold2 seem to be relevant for IDPs [5]. Firstly, AlphaFold2 predicts well regions that will not fold under any normal circumstances [6]. Secondly, it can predict segments that might fold upon binding to other molecules, *i.e*. the predicted structure is that of the protein in complexed form with other proteins or lipid membranes. This context-dependent folding has been predicted by other bioinformatics tools before [7, 8], allowing to detect functional regions in IDPs that interact with other molecules.

IDPs or IDRs do not fold into a stable 3D structure, but rather exist as an ensemble of conformations. Their conformational properties depend on their amino acid composition, and upon molecular interactions, secondary structures do form. IDPs are, hence, physicochemically very different from denatured globular proteins [9]. Due to their specific properties as polymeric chains, several biological functions have been attributed to IDPs and IDRs. These include, but are not limited to, acting as molecular rulers, forming membraneless organelles, protecting from collapse under plant dehydration [10, 11], increasing the avidity of clamp binders [12], and binding to lipid membranes [13]. Conformational plasticity and the ability for context-dependent folding are central for such functions of IDPs.

The increase in nerve conduction velocity, enabled by the myelin sheath, is mandatory for the normal functioning of the vertebrate nervous system. Myelin is a multilayered proteolipid membrane in the central and peripheral nervous system (CNS and PNS, respectively), which is formed by myelinating glia and wraps around selected axons. The compacted myelin membrane carries a unique set of proteins, which are either integral or peripheral membrane proteins binding two or more lipid bilayers together into multilayers. Myelinating cells express several specific IDPs, which are crucial for the correct formation and stability of the myelin membrane multilayer [14]. Such proteins include the IDPs myelin basic protein (MBP) and periaxin, and the cytoplasmic IDR of myelin protein zero (P0ct). The folding of disordered myelin proteins has been studied both with the full proteins and with selected peptide segments [15–21], allowing detection of membrane interaction sites and lipid binding-induced folding into helices.

Intriguingly, many mutations linked to peripheral human neuropathies, mainly different forms of Charcot-Marie-Tooth disease (CMT), are found in IDPs or IDRs in myelin proteins. This highlights the important functional/structural role of these protein segments, whereby they may be important membrane interaction sites or participate in protein-protein complexes. CMT mutations are found in both P0ct [22–27] and in extended disordered regions of periaxin [28–31]. Puzzlingly, thus far, no mutations in MBP have been linked to any disease, despite its high abundance and apparent importance for myelin structure.

The high-resolution 3D structure determination of MBP and P0ct has proven to be difficult, if not impossible, given the futile attempts over the past three decades. Here, we extracted information through the most recent computational predictions, namely, the AlphaFold2 models of MBP and P0ct. For both proteins, we analyse earlier experimental data in light of the AlphaFold2 models and show that AlphaFold2 models are valuable even in the case of highly flexible, disordered proteins, helping to understand the function and interactions of these proteins at the molecular level.

## Methods

### Generation of molecular models for MBP and P0ct

AlphaFold2 [1] was run on the Google ColabFold site [32], giving as input only the amino acid sequence of mouse 18.5-kDa MBP isoform and human P0ct. The resulting 5 models were all relaxed with the Amber implementation in AlphaFold2. The models were used as such for further analyses.

### SAXS data analysis

SAXS data for mouse MBP and human P0ct from our earlier publications [13, 33] were directly used to assess the fits of the AlphaFold2 models to the experimental data. Specifically, the programs CRYSOL [34], OLIGOMER [35], and EOM [36] were used. R_g_ values were additionally estimated using the Guinier plot in PRIMUS [35] and with the Debye formalism, as described [37, 38]. D_max_ was manually estimated using GNOM [39], such that the distance distribution had a reasonable shape and the fit to the raw SAXS data was optimal.

### Bioinformatics and structure analysis

Sequence-based predictions for both proteins have been published [40, 41]. The highest-scoring AlphaFold2 models of MBP and P0ct were docked onto planar and curved lipid bilayers with properties of a eukaryotic plasma membrane, using the PPM server [42]. Visualisation and surface electrostatics calculations were carried out in PyMOL [43] with the APBS [44] plugin. Previously published bioinformatics and structural results were considered and further analysed in relation to the current predictions.

## Results and Discussion

The myelin sheath is a multilayered proteolipid membrane with unique proteins, such as myelin basic protein (MBP), myelin protein zero (P0) and peripheral myelin protein 2 (P2), which all function in the tight stacking of the lipid bilayers, ensuring correct functioning of the myelin sheath. Malfunction of this system, for example mutations in myelin proteins or autoimmune attack, results in neurological diseases, such as multiple sclerosis, various types of Charcot-Marie-Tooth disease (CMT), Dejerine-Sottas syndrome, and congenital hypomyelination. While some myelin proteins are structurally well-defined, like P2, there several myelin IDPs that only change their conformation upon different interactions, including MBP and P0ct.

Encouraged by work from others on understanding IDPs and their conditionally folded segments [45–47], we set out to analyse earlier experimental SAXS data [13, 33] as well as literature in light of AlphaFold2 models of MBP and P0ct. We expected to get an improved picture about the folding of these two myelin-specific proteins, when they interact with membrane surfaces, since we expected AlphaFold2 to predict ordered conformations with high confidence. While AlphaFold2 cannot predict accurate atomistic 3D structures for IDPs, and we did not expect high-resolution information on folding, we focused on observing if we can better explain the unique membrane interactions of these two myelin IDPs. Furthermore, AlphaFold2 can predict, which parts of the IDP do not fold upon interactions with other molecules or surfaces.

For both MBP and P0ct, data from various biophysical experiments have been published [33, 41, 48–53], showing folding of both proteins upon membrane binding, while they remain unfolded in aqueous solution. The regions binding to membranes have been mapped to specific segments, mainly those prone to fold into helices according to predictions. Several studies have shed light on more details of membrane binding by focusing on peptides corresponding to the membrane-binding sites. The relation of the membrane-binding sites of myelin proteins with possible autoantigenic epitopes in disease [15, 40], such as multiple sclerosis, together with molecular mimicry of certain viruses like EBV [54], suggests that detailed fundamental studies on myelin protein membrane interactions can give new insights into both myelin biology and pathology. How useful might an AI-based prediction of protein 3D structure be in this scenario?

### Myelin basic protein – the molecular glue in myelin

MBP lacks a well-defined structure in aqueous solution, but it changes conformation depending on its interactions and the chemical environment. In fact, its disordered nature was described already nearly 50 years ago, well before IDPs had become a central topic in protein biochemistry [55]. MBP is one of the best-characterized proteins of the myelin sheath, playing a role in many different interactions, oligodendrocyte proliferation, stabilization of myelinogenesis and membrane stacking [56]. The oligodendrocyte lineage (golli) gene gives rise to a variety of MBP isoforms ranging from 14.0 kDa to 21.5 kDa, with the 18.5 kDa isoform being the most predominant one in human mature myelin [57]. Furthermore, all isoforms of MBP have a positive charge depending on post-translational modifications (PTMs), referred to as C1 to C8, where C1 is the most basic isomer (net charge of +19 at physiological pH), with the least amount of PTMs [58]. The most common PTMs are phosphorylation and citrullination, with the C8 isoform being the least basic isomer having decreased ligand interactions compared to the C1 isoform [59].

Despite early support for a β-sheet MBP structure, Mendz et al. suggested in 1990 that certain interaction sites within the molecule form helices when mixed with detergent micelles [60]. Especially this model has been considered for the central helical segment between residues 82 and 93 in mouse 18.5-kDa MBP (85-96 in human MBP), and an α-helical model would facilitate interactions with lipid head groups [61]. With the use of electron paramagnetic resonance spectroscopy and molecular dynamics simulations, later studies indicated an a-helical structure for this segment [62]. The depth profile indicated an amphipathic α-helix penetrating up to 12 Å into the myelin-like membrane. MBP has high levels of arginine and low levels of glutamatic acid, which contributes to its basicity, required for its interaction with negatively charged phospholipids. With the less basic isoform C8, the C-terminal region dissociated from the membrane, while the N-terminal site was more mobile than for C1 [52]. In the same study, the Phe-86/Phe-87 motif was important for the formation of the helix and its attachment to lipids [52, 59]. Overall, three MBP segments, T33-D46, V83-T92 and Y142-L154, have been experimentally found to be a-helical, located close to the N and C terminus and in the central region of MBP; the formation of these helices is regulated by the local hydrophobic interactions between the nonpolar surface of the helix and the lipid bilayer [57]. Wang et al. further confirmed with SPR that the interactions between MBP and lipid monolayers are electrostatic, and the protein binds strongly with increased fraction of negative headgroups [63].

The most abundant and experimentally best studied isoform of MBP, the 169-residue 18.5-kDa isoform, was used for the analyses here. The five models of MBP predicted by AlphaFold2 are shown in **Fig. 1A**. All 5 models have similar folds and dimensions. The superposition of the obtained models creates a structural ensemble akin to those obtained *e.g*. from EOM based on SAXS data [ref], but in this case, the models are based on sequence alone.

**Figure 1.**
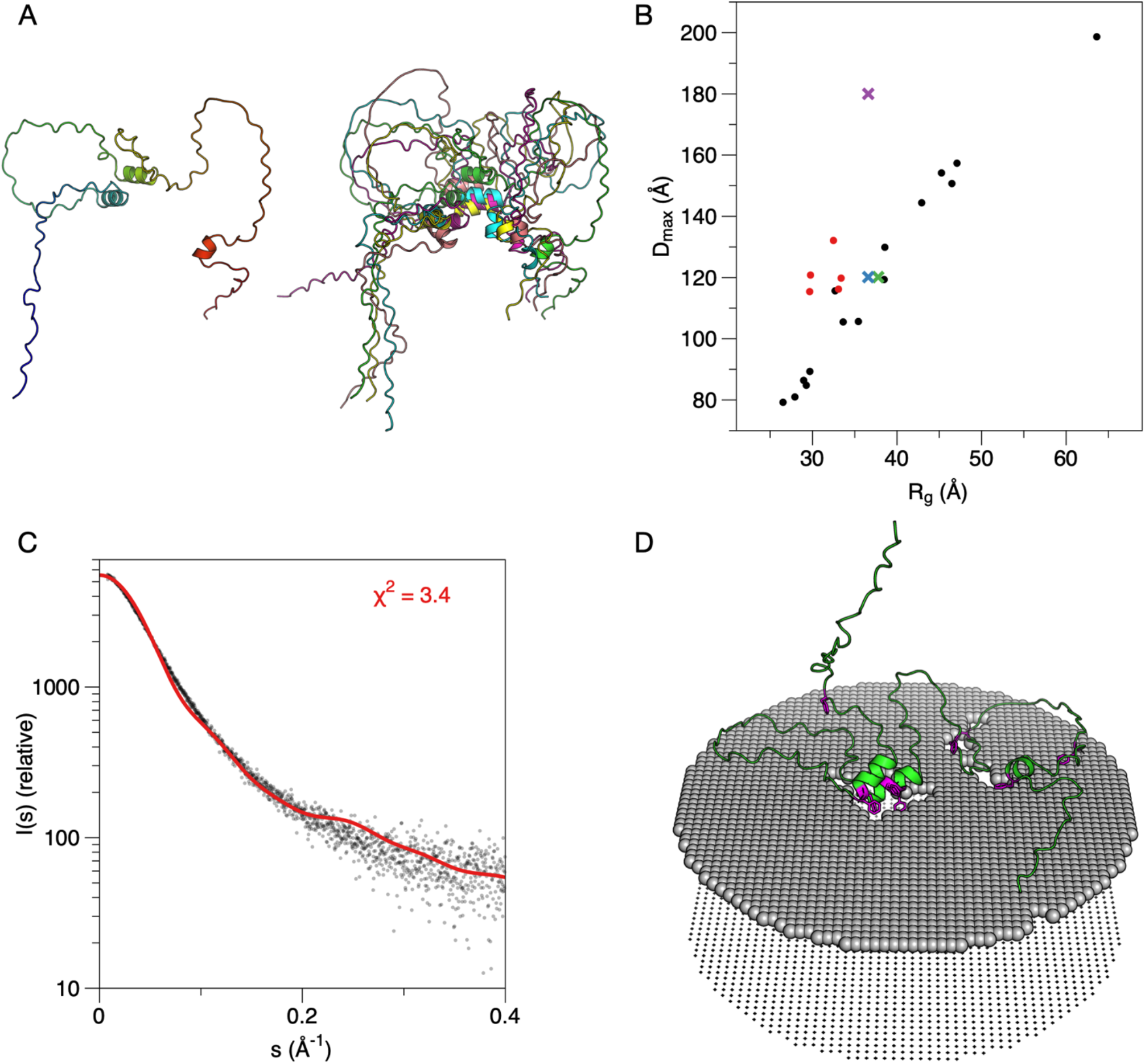
AlphaFold2 models of MBP. A. The top ranked (left) model and superposition of the 5 obtained models (right). Note how the overall dimensions and shape are similar, and that the three short helices cluster in the middle in all cases. B. Comparison of the AlphaFold2 models (red dots) and the EOM ensemble based on solution SAXS data (black dots). The average R_g_/D_max_ from EOM is shown with a green cross, the Guinier R_g_ with D_max_ from EOM with a blue cross, and the Guinier R_g_ with manually determined D_max_ from GNOM with a magenta cross. While the models cluster close to the average experimental values from EOM, they systematically have a lower R_g_, which is a sign of the presence of folded structure. C. Fit of the top ranked model alone to experimental SAXS data [13]. A similar overall shape is obvious, and the fluctuations are related to the secondary structures in the model and their clustering. D. Docking of the top ranked model onto a lipid bilayer surface suggests membrane interactions by the helices and lends support to the hypothesis that the predicted model reflects the membrane-bound state. Phe residues are highlighted in magenta.

A more detailed analysis of the predicted structures is, therefore, warranted. A comparison of their R_g_ and D_max_ to those obtained from EOM is shown in **Fig. 1B**. The AlphaFold2 models apparently have a smaller R_g_ for the same D_max_, when compared to the flexible random-chain models produced by EOM – this reflects the presence of folded secondary structure elements in the models; also note how the predicted helices tend to cluster together in the models (**Fig. 1A**). Three helices are predicted in all models (at residue ranges 36-45, 83-92, and 148-153), and these correspond to the membrane- and calmodulin-binding sites identified earlier [18–20, 59] that become α-helical upon binding. This observation indicates that AlphaFold2 has predicted, at least partially, the membrane-bound conformation of MBP, rather than the form free in solution.

The highest-ranked AlphaFold2 model of MBP provided a reasonable fit alone to the raw experimental SAXS data (**Fig. 1C**). The fit was not improved by fitting all five models simultaneously using OLIGOMER. This does indicate that the top model alone is a reasonable representation of MBP structure – with the caveat that we expect AlphaFold2 to rather predict the membrane-bound model than the one free in solution. With this in mind, we modelled the AlphaFold2 structure onto a membrane surface (**Fig. 1D**). Whether the MBP helices cluster together on the membrane, as seen in the AlphaFold2 models, is not known. Notably, the two main helices that come together in the models both harbour a double Phe motif, which has been shown to be crucial for MBP function in membrane stacking [64]. MBP compacts drastically upon being embedded between two membranes [13], which are only 3 nm apart, and this clustering of helical segments could be a mechanism of structural compaction. Given that the internal helices of MBP would cluster together as predicted by AlphaFold2, this could represent the intermolecular mechanism that ultimately results in the liquid-liquid phase separation of MBP [64]. In addition, the conformation could coarsely represent the formation of a gel-like protein phase on the surface of a lipid bilayer, which we earlier observed using cryo-EM and neutron reflectometry [13].

### The cytoplasmic domain of P0 – similar but different to MBP

P0 is the major protein in the PNS myelin, being primarily expressed in Schwann cells. The Ig-like extracellular domain is the only part of the P0 that has been structurally characterized at atomic detail using X-ray crystallography [65, 66]. The transmembrane domain of full-length P0 contains a single α-helix. Within the myelin sheath, the P0 is assumed to form homodimers due to the “glycine zipper” associated with the domain [67]. P0 molecules are believed to oligomerize between proteins located on apposing membranes [68], *via* both the extracellular and intracellular domains. The cytoplasmic domain P0ct is not only important for membrane stacking at the PNS major dense line, but given its nature as part of a transmembrane protein, it could be involved in P0 trafficking, which further is regulated by post-translational modifications in the P0ct [51].

P0ct is comprised of 69 residues, being disordered in solution and having a high positive charge. However, CMT disease-causing mutations have been identified within this domain [22–27], highlighting its importance for proper myelination. Like MBP, P0ct folds into helical structures upon interactions with lipid membranes [33, 41]. P0ct has a strong (+15) positive charge, carrying 21 basic and 6 acidic residues [69]. Full-length P0 contains three cysteine residues; Cys21 and Cys98 form the disulphide bond of the extracellular domain. The third cysteine resides in the cytoplasmic domain at the junction between the transmembrane domain and the cytoplasmic tail [70]. This conserved cysteine is an acylation site and often undergoes palmitoylation [70, 71]. A mutation of this residue results in loss in both the attachment of fatty acid and the adhesiveness of P0 [72]. When mutated or truncated, the ability of P0ct to hold two membranes together is lost, suggesting that acylation participates in myelin stability [72, 73]. P0ct as a free peptide in aqueous solution is unfolded, as determined by CD spectroscopy, but gains secondary structure upon lipid interactions. The folding was earlier suggested to be mostly ß-sheets, but later studies strongly support a more α-helical conformation [33, 48]. We showed that full-length P0 organises into dimers in a zipper-like way when reconstituted into small unilamellar vesicles, and P0ct in a lipidic environment induced Bragg peaks when subjected to small-angle X-ray diffraction in a concentration-dependent manner [33], indicating spontaneous assembly of ordered semi-crystalline structures.

P0ct models are shown in **Fig. 2A**, and their R_g_ and D_max_ distribution with respect to EOM results are shown in **Fig. 2B**. The outcome is similar to MBP, giving a further indication of the shared physicochemical properties between MBP and P0ct. One helix is predicted at the beginning of the P0ct; this segment is expected to bind along the membrane surface, and to be anchored to the membrane tightly *via* both the transmembrane domain and the palmitoylated Cys [21]. A second helix is in the middle region of P0ct and represents an additional membrane anchor [41]; whether it binds to the same or the apposing membrane in myelin, is currently not known. Mutations D224Y and R227S at this helical site are linked to CMT [22–25, 33, 41]. Intriguingly, this helical site is also a hotspot for PTMs, such as phosphorylation, that have been linked to the trafficking of P0 during myelination [51].

**Figure 2.**
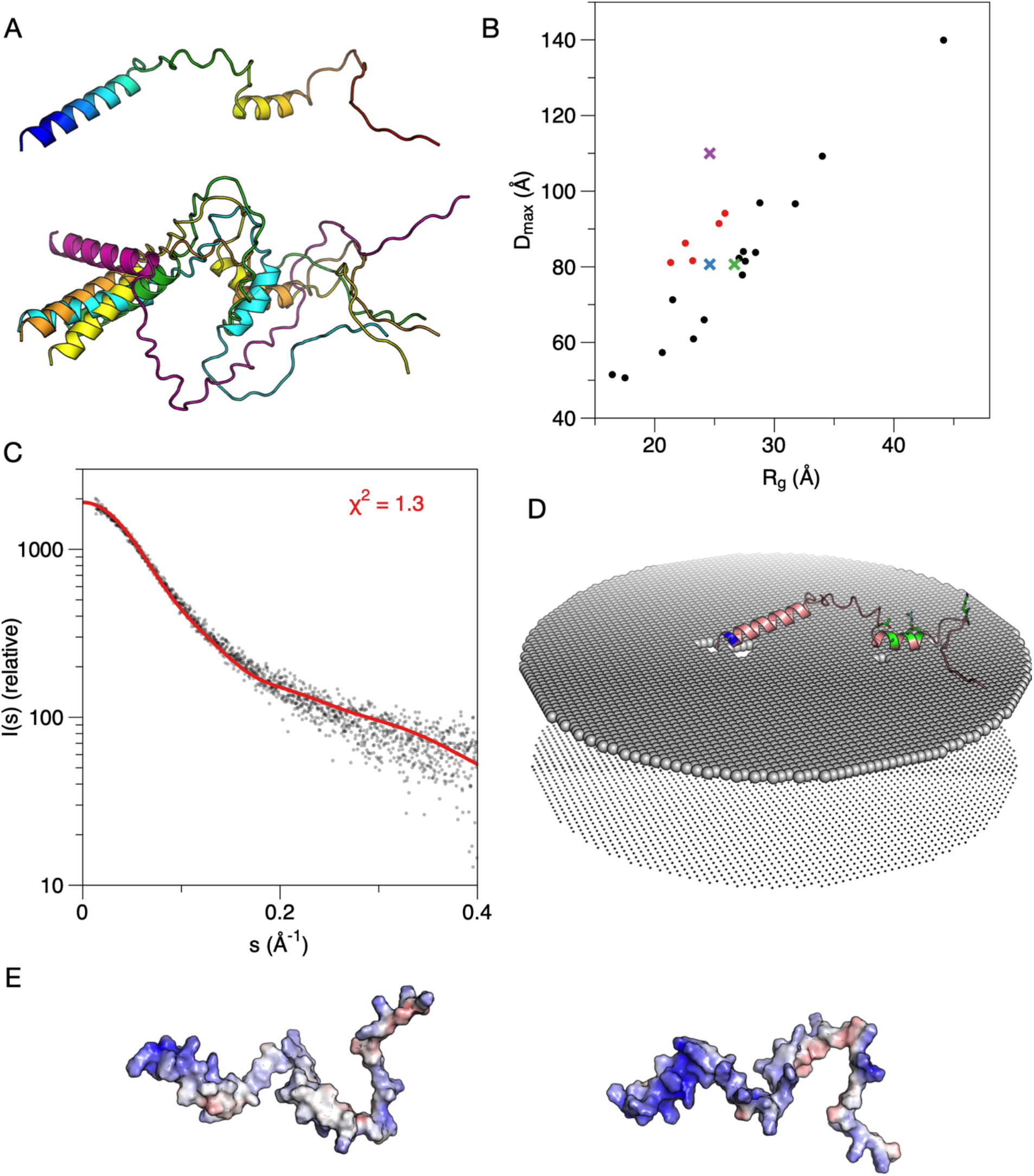
AlphaFold2 analysis of P0ct. A. The top-ranked model (top) and all five models superimposed (bottom). All models include two helices and have similar dimensions. B. Comparison of the P0ct AlphaFold2 models (red dots) and the full EOM ensemble (black dots). The green cross indicates the average values from EOM analysis of experimental data [33]. The Guinier R_g_ with D_max_ from EOM is marked with a blue cross and the Guinier R_g_ with manually determined D_max_ from GNOM with a magenta cross. C. The top ranked P0ct model fits the raw SAXS data very well. D. Docking of the P0ct model onto a membrane surface. Blue indicates the location of the Cys residue close to the transmembrane domain, and CMT mutation sites are coloured green. E. Electrostatic surface of P0ct from two orientations. The face binding the membrane is hydrophobic (left), while the opposite side is positively charged (right).

The highest-ranked AlphaFold2 model of P0ct fits well to the SAXS data (**Fig. 2C**). OLIGOMER fitting of the 5 top models did not improve this fit, and the full EOM ensemble fit only slightly better than the single predicted model, indicating that the model reproduces well the average size and shape of P0ct in solution. Furthermore, the original *ab initio* model built based on the SAXS data [33] again only provides a marginally better fit than the AlphaFold2 model (**Table 1**).

**Table 1.** Fits of different models to experimental synchrotron SAXS data. The values given in the table are χ^2^ for the fit between model and experimental data.

| Protein | MBP | P0ct |
| --- | --- | --- |
| Highest-ranked AlphaFold2 model alone | 3.4 | 1.3 |
| OLIGOMER solution, fitting all 5 AlphaFold2 models | 4.3 | 1.3 |
| Full EOM ensemble | 1.0 | 1.0 |
| Chain-like *ab initio* model (GASBOR) | 1.1 [13] | 1.1 [33] |

The helices predicted by AlphaFold2 on P0ct coincide with earlier identified functional segments interacting with lipid membranes (see above). Furthermore, peptides encompassing both of the predicted helices in P0ct have been used to generate animal models of human autoimmune neuropathies [74]. Hence, these sites are known to show disorder-to-order transitions and either carry point mutations for CMT disease or autoantigenic epitopes that induce experimental autoimmune neuritis and possibly human Guillain-Barré syndrome [41, 75].

Interestingly for an IDP, a total of 6 missense mutations have been identified in the P0 cytoplasmic tail, linked to human CMT [22–27]. The location of these mutations in the model is depicted in **Fig. 2D**, which also shows the predicted orientation of the P0ct model on a membrane surface. Importantly, these mutations are concentrated within the central region of P0ct, mainly in the membrane-binding helix. Intriguingly, one of them, D224Y, causes both hypermyelination in patients and increased membrane stacking *in vitro* [24, 41]. The model suggests the CMT mutations in P0ct could directly affect its membrane interactions in the tightly confined space of the myelin major dense line. The electrostatic potential surface of the P0ct highest ranked model is shown in **Fig. 2E**, indicating a positively charged face and a hydrophobic surface, compatible with amphipathic membrane interactions. Considering the tightly confined space of the PNS major dense line, we currently cannot be sure whether the middle helical segment of P0ct binds to the same membrane as the transmembrane domain, or if it reaches over and inserts itself into the apposing cytoplasmic leaflet.

### Additional notes on fitting to SAXS data

While for both MBP and P0ct, EOM gives the best fit to the experimental SAXS data, it is quite remarkable how well the single first-ranked AlphaFold2-based IDP models of MBP and P0ct fit the SAXS data published before. Single *ab initio* models fit the data slightly worse than the full conformational EOM ensembles, showing the presence of several conformations. Especially in the case of P0ct, a single AlphaFold2 model fits very well to the solution SAXS data from recombinant P0ct. IDPs are often not straightforward cases for SAXS studies, as discussed in recent literature [76]. The data do indicate that single models of IDPs from AlphaFold2 can complement SAXS data and provide reliable representations of the IDP at low resolution. As for folded proteins, therefore, such models can be valuable additions to support experimental data and help in setting up and evaluating hypotheses on structure-function relationships.

In essence, for both IDPs studied here, the AlphaFold2 models are close to the average D_max_ of the disordered EOM ensemble and the R_g_ obtained from Guinier plot (**Fig. 1B, 2B**). On the other hand, D_max_ determined in a traditional way subjectively from distance distribution more estimates the absolute largest D_max_ in the population instead of the average (**Fig. 1B, 2B**). For IDPs, Debye formalism provides a more relevant R_g_ than the Guinier plot [37], and indeed, this value is close to that of the EOM ensemble average R_g_. From Debye analysis, the R_g_ for MBP is 42.1 Å and that for P0ct 26.2 nm. These analyses further indicate that the AlphaFold2 models do not represent the disordered ensembles in solution, but slightly compacted conformations, possibly corresponding to the lipid-bound conformation.

### Conclusions

The intermembrane compartment harbouring MBP and P0ct in the PNS, the major dense line, is very tight, with a spacing of only ~3 nm between the bilayers. This indicates, together with the expected molecular dimensions of both MBP and P0ct, that both proteins must interact with two membranes simultaneously. These interactions are enabled by both the membrane anchor segments forming α-helices as well as the flexible, disordered segments between them. The use of AlphaFold2 models in this short report has highlighted that molecular models can be used to obtain additional details of functional significance in combination with earlier and current experimental data. In some cases, conclusions can be drawn, for example, on the effects of disease mutations on IDP structure and interactions. While the overall 3D structure of an AlphaFold2 model of an IDP will not be accurate, nor does it give much information about conformational ensembles, it does give relevant information about average molecular size and shape, as well as segments that are likely to fold into secondary structure upon molecular interactions. Accordingly, it has not escaped our attention that for both IDPs studied here, the highest-ranked AlphaFold2 model fits the solution SAXS data remarkably well, considering the only input to modelling was the sequence of an IDP. Hence, AlphaFold2 does provide meaningful information on at least the overall size and shape of these IDPs, but it additionally has the power to predict interaction sites and conditionally folded segments linked to them. Hence, in combination with experimental biophysical and structural work on IDPs, the predicted models can help explain molecular mechanisms in IDP biology and disease.

